# Musashi-2 (MSI2) regulation of DNA damage response in lung cancer

**DOI:** 10.1101/2023.06.13.544756

**Authors:** Igor Bychkov, Alexander Deneka, Iuliia Topchu, Rajendra P. Pangeni, Christopher Lengner, John Karanicolas, Erica A. Golemis, Petr Makhov, Yanis Boumber

## Abstract

Lung cancer is one of the most common types of cancers worldwide. Non-small cell lung cancer (NSCLC), typically caused by *KRAS* and *TP53* driver mutations, represents the majority of all new lung cancer diagnoses. Overexpression of the RNA-binding protein (RBP) Musashi-2 (MSI2) has been associated with NSCLC progression. To investigate the role of MSI2 in NSCLC development, we compared the tumorigenesis in mice with lung-specific *Kras*-activating mutation and *Trp53* deletion, with and without *Msi2* deletion (KP versus KPM2 mice). KPM2 mice showed decreased lung tumorigenesis in comparison with KP mice what supports published data. In addition, using cell lines from KP and KPM2 tumors, and human NSCLC cell lines, we found that MSI2 directly binds *ATM/Atm* mRNA and regulates its translation. MSI2 depletion impaired DNA damage response (DDR) signaling and sensitized human and murine NSCLC cells to treatment with PARP inhibitors *in vitro* and *in vivo*. Taken together, we conclude that MSI2 supports lung tumorigenesis, in part, by direct positive regulation of ATM protein expression and DDR. This adds the knowledge of MSI2 function in lung cancer development. Targeting MSI2 may be a promising strategy to treat lung cancer.

**Significance:** This study shows the novel role of Musashi-2 as regulator of ATM expression and DDR in lung cancer.

## Introduction

Lung cancer is one of the most frequently diagnosed cancers and is the leading cause of cancer related death worldwide. Based on World Health Organization data (1), lung cancer counts for >2 million (11.4%) of new cases and 1.8 million (18%) of lung cancer deaths in 2020. Owing to the absence of clinical symptoms in early-stage disease and limitations of effective screening programs, most lung cancers are diagnosed in advanced stages. The most common type of lung cancer is non-small cell lung cancer (NSCLC), representing 85% of total lung cancer cases (2), which includes adenocarcinoma, squamous cell carcinoma and large-cell carcinoma histologic subtypes. Recent cancer genome sequencing efforts have defined the complex of genomic aberrations that lead to the lung cancer development (3). In NSCLC, druggable alterations include mutations in Epidermal Growth Factor Receptor (EGFR), ALK and ROS rearrangements and others, and occur in ∼10-60% of tumors (4). Approximately 30% of lung adenocarcinomas are driven by KRAS mutations (5), some of which are druggable. However, many NSCLC do not respond well to treatment, and overall survival for NSCLC patients diagnosed at a late stage remains 26%(6).

KRAS mutations often co-exist with mutations impairing activity of the *TP53* tumor suppressor, contributing to increased genomic damage (7). The DNA damage response (DDR) signaling network regulates activation of transcription, cell cycle, apoptosis, senescence, and DNA repair processes in response to DNA damage (8). Coordination of these biological processes is critical for cell survival when DNA replication is perturbed or when cells are treated with mutation-inducing agents. The core of the DDR mechanism is a pair of related protein kinases, Ataxia-telangiectasia mutated (ATM) and Ataxia-telangiectasia and Rad3-related protein (ATR), which are activated by DNA damage. ATM is regulated by the MRN (Mre11-Rad50-NBS1) complex which senses double-strand breaks (DSBs), whereas ATR is regulated by ATRIP (ATR-interacting protein), which senses single-strand DNA (ssDNA) generated by processing of DSBs, as well as ssDNA present at stalled replication forks (9-12). The poly (ADP-ribose) polymerase (PARP) protein is critical for several forms of DNA repair, including nucleotide excision and base excision repair (NER and BER) and homologous recombination (HR), and supports repair of DNA damage induced by alkylating agents and chemotherapy. Several recent clinical studies have recently shown potential efficacy of PARP inhibitors in NSCLC, especially when combined with platinum-based chemotherapy or immunotherapy, although there is lack of clarity of specific biomarkers for patient selection that benefit from this approach (13-16). Interestingly, up to 40% of human lung adenocarcinomas lack ATM protein expression, again implicating altered DDR in a subset of NSCLC (17). This provides the basis for a potentially therapeutic approach, as recent publications show that ATM deficiency in NSCLC is associated with higher sensitivity to the PARP inhibitor olaparib and the ATR inhibitor AZD6738 (18-20).

Our previous work has shown the elevated expression of MSI2 in advanced NSCLC (21,22). MSI2 is an RNA-binding protein which regulates the stability and translation of target mRNAs via recognition of specific core motifs in the 3’-untranslated region (UTR) (23-25). MSI2 and its homolog, MSI1, regulate multiple critical biological processes relevant to stem cell compartment maintenance, cancer progression, and cancer drug resistance (26,27), and have elevated expression in numerous types of cancer (21,24,28-31). In studies of cell lines established from highly metastatic tumors arising in mice expressing lung-specific activated KRAS and TP53 deletion (129S/Sv-*Kras*^*tm3Tyj*^/J;*Trp53*^*t*m1Brn^/J (KP) mice), we previously established MSI2 as upregulated in a subset of aggressive NSCLC, and demonstrated a specific role for MSI2 in promoting metastasis in these tumors, through induction of TGFβR1 and its effector SMAD3 (22). In other studies, using human cell models, we have shown that MSI2 directly binds to EGFR mRNA and regulates EGFR protein expression, while depletion of MSI2 has led to increased sensitivity to EGFR inhibitors in EGFR-mutant NSCLC (21). Also, we have shown that MSI2 directly regulates PTEN and VEGFR2 protein levels in NSCLC (32). These data suggested a broad role for MSI2 in increasing tumor aggressiveness.

In this study, we analyzed the effect of simultaneous deletion of *Msi2* (33) on the pathogenesis and therapeutic response of KP mice (34), creating and analyzing a new (*Kras*^*mut*^*/P53*^*KO*^*/MSI2*^*KO*^ or KPM2 model. The new murine KPM2 model showed decreased total tumor number and burden in comparison with positive control (KP) and supports previously published papers(21,22,32,35,36) Furthermore, murine lung cancer cell lines, which were generated from those models showed decreased proliferation activity of KPM2 cells in comparison with KP. Subsequent experiments showed that MSI2 affects DDR, as it directly regulates ATM protein expression in murine and human NSCLC. Also, MSI2 deficiency leads to higher sensitivity to PARP (Poly (ADP-ribose) polymerase) inhibitor in murine and human NSCLC.

## Materials and methods

### Animals

All animal experiments were approved by the Institutional Animal Care Committees at Fox Chase Cancer Center and Robert H Lurie Comprehensive Cancer Center. To generate A novel KP-MSI2 model (129S/Sv-*Kras*^*tm3Tyj*/J^;*Trp53*^*t*m1Brn^/^J^; *Msi2*^*-/-*^), we crossed 129S/Sv-*Kras*^*tm3Tyj*/J^;*Trp53*^*t*m1Brn/J^ (KP) mice(34) with C57BL/6 floxed *Msi2*^*-/-*^ *Cre* transgenic mice (33). Following lung-specific induction with Adeno-Cre, each 6 week old mouse get 2.5 ×10^6^ units of Cre recombinase adenovirus (Vector Biolabs, Malvern, PA), the 129S/Sv-*KP* mice develop lung disease similar to that of humans; large papillary adenomas are seen after 6 weeks and adenocarcinoma after ∼10 weeks, in 4 month old animals (34). Mice were euthanized and their lungs, heart, liver, spleen were collected for paraffin blocks, and lung cancer cell lines were established from fresh lung mouse tumors, also ∼5 months after activation, Cre induction in the lungs and euthanasia of the mice.

### Mouse genotyping

Small segments of mouse tails (0.5∼1.0 cm) were collected and digested in 150 μL of DNA-lysis buffer with 0.2 mg/ml of proteinase K for overnight at 56 ºC. Then, DNA samples were diluted in 50 times and mixed with DreamTaq master mix (2X) (Thermofisher) following vendor protocol for genotyping PCR. Then, DNA electrophoresis was performed in 2% agarose gel. Bands detection was performed using imaging system ChemiDoc (Bio-Rad). All primers used in this procedure are noted in Supp Table S1.

### Assessment of *in vivo* tumor growth

For *in vivo* studies 2 × 10^6^ of KP-1 and KPM2-2 cells were injected subcutaneously (s.c.) in the right flank of 6 week old mice C57BL/6 (total: 20 males, 20 females) from The Jackson Laboratory (ME, USA). All mice were over 18g at the start of live-phase study. When tumor volumes reached 100 mm^3^ animals were randomly assigned to the control or experimental groups (n = 10 mice/group, 5 males, 5 females). The mice were treated with 5% DMSO, 40% PEG 300, 5% Tween-80, 50% sterile H_2_O (vehicle) or olaparib 50 mg/kg in 5% DMSO, 40% PEG 300, 5% Tween-80, 50% sterile H_2_O. Treatment was perfomed by daily oral gavage. Tumors were measured twice weekly, and their volumes were calculated with the formula: [volume = 0.52 × (width)^2^ × length]. Mice were euthanized when tumor volume exceeded 1500 mm^3^ or on 31^st^ day of experiment.

### Immunohistochemistry of NSCLC samples

Scientific review committee (SRC) and IRB approvals were obtained for de-identified samples from Fox Chase Cancer Center, both at Fox Chase Cancer Center and at Robert H Lurie Comprehensive Cancer Center. Tissue samples were stained for MSI2, MSI1, ATM and CyclinB1 via immunohistochemical (IHC) approach and hematoxylin and eosin (H&E) stained sections were used for morphological evaluation purposes, and unstained sections were used for IHC staining using standard methods. Briefly, 5 μm formalin-fixed, paraffin-embedded sections were deparaffinized and hydrated. Sections were then subjected to heat-induced epitope retrieval with 0.01M citrate buffer (pH 6.0) (MSI2, MSI1, ATM) or EDTA buffer (CyclinB1). Endogenous peroxidases were quenched by the immersion of the slides in 3% H_2_O_2_ solution. The sections were incubated overnight with primary antibodies at 4°C in a humidified slide chamber. As a negative control, the primary antibody was replaced with normal mouse/rabbit IgG to confirm absence of specific staining. Immunodetection was performed using the Dako Envision+ polymer system and immunostaining was visualized with the chromogen 3, 3′-diaminobenzidine. All slides were viewed with a Nikon Eclipse 50i microscope and photo-micrographs were taken with an attached Nikon DS-Fi1 camera (Melville, NY, USA). For IHC quantification, each spot was examined by board-certified pathologists (ED and NK) who assigned a score of 0 (no staining), 1+ (weak staining), 2+ (moderate staining), and 3+ (strong staining) within carcinomatous areas. The score for each of the two tumor spots was averaged for statistical analysis. The H-score, which ranges from 0 to 300, was calculated using the following formula: [1x(% cells 1+) + 2x(% cells 2+) + 3x(% cells 3+)], which reflects staining intensity as well as percentage of positive cells. A sum of p2 and p3 represents a sum of 2+ and 3+ cells (2x (% cells 2+) + 3x(% cells 3+), which excludes 1+ cells. Clinical information (Supp Table S5) from the repository database was abstracted in an anonymized fashion.

### Establishment of cell lines

Murine lung tumors were collected and used to generate. murine NSCLC cell lines by standard approaches. Briefly, the lines 129S/Sv-*Kras*^*tm3Tyj*/J^;*Trp53*^*t*m1Brn^/^J^; *Msi2*^*-/-*^ (KPM2-1, KPM2-2, KPM2-3) and *Kras*^*tm3Tyj*/J^;*Trp53*^*t*m1Brn/J^ (KP-1, KP-2, KP-3) were prepared from freshly isolated cancer cells. Initial stocks were cryopreserved, and at every 6-month interval, a fresh aliquot of frozen cells was used for the experiments. The tumor pieces were incubated with collagenase (400 u/ml) at 37 ºC for 2 hours, then we used cell strainer (70 μm) for cells separation. After that, the cells were incubated with ACK lysis buffer at room temperature for 1-2 minutes, then the cells were washed and transferred to T25 flask with B27 cell culture media for propagation.

### Cell culture of human cancer cell lines

Human lung cancer cell lines with KRAS-mutations (A549 and H441) were obtained from the American Type Culture Collection (ATCC). No additional authentication was performed. All cells were cultured in RPMI 1640 (Gibco, Gaithersburg, MD) supplemented with 10% FBS (Hyclone, Logan, UT), penicillin (100U/ml), streptomycin(100μg/ml), sodium pyruvate (1 mM) and non-essential amino acids (0.1 mM) under conditions indicated in the figure legends.

### Antibodies and drugs

Anti-MSI2 (#ab76148), anti-MSI1 (#ab21628), anti-phCHK2 T68 (#ab278548) were obtained from Abcam (Cambridge, UK). Anti-phATM S1981 (#GTX132146), anti-ATM (#GTX70103), anti-ATR (#GTX128146), anti-phCDC25A (#GTX55131), anti-CDC25A (#GTX102308) were obtained from GeneTex, Inc. (Irvine, CA). Anti-phH2A.X S139 (#9718), anti-ATM (#2873), anti-CHK2 (#2662), anti-phCHK1 S345 (#2348), anti-CHK1 (#2360), anti-β-actin (#3700), anti-rabbit HRP-linked (#7074), anti-mouse HRP-linked (#7076) were obtained from Cell signaling (Danvers, MA). Anti-MSI1 (#AF2628) was obtained from R&D systems (Minneapolis, MN) Anti-ATM (#27156-1-AP) was obtained from Proteintech (Rosemont, IL). Olaparib (#HY-10162), cisplatin (#HY-17395), doxycycline (#HY-N0565) were obtained from MedChemExpress (Monmouth Junction, NJ).

### Vector construction and lentivirus production

To generate stable human cell lines with inducible MSI2 knockdowns, self-complementary single-stranded DNA oligos (Supp Table S2) were annealed and cloned into AgeI/EcoR1 sites of Tet-pLKO-puro vector (Addgene plasmid # 21915). All constructs were validated by direct sequencing. All generated cell lines used in the study are noted in Supp Table S3.

### Western blot analysis

Cell lysates preparation and Western blot analysis were performed using standard methods as previously described(21). Signals were detected by X-ray films and digitized by photo scanner. Image analysis was done using ImageJ (version 1.53e, National Institutes of Health, Bethesda, MD), with signal intensity normalized to β-actin; 3-4 repeats were used for each quantitative analysis. Final data was analyzed in GraphPad Prism by unpaired t-test to determine statistical significance.

### Reverse transcription and qPCR

RNA was extracted using a phenol-chloroform based method. RNA concentration and quantity were measured using NanoDrop Lite (cat# ND-LITE ThermoFisher Scientific). First strand cDNA synthesis was performed with iScript cDNA synthesis kit (cat#1708841, Biorad, California, USA) according to manufacturer’s instructions. The generated cDNA was diluted tenfold and used as a template for qPCR, which was performed with Applied Biosystems QuantStudio 3 system using PowerTrack™ SYBR Green Master Mix (Applied Biosystems). Relative quantification of genes expression was performed using 2^-ΔΔCt^ method, using primers indicated in Supp Table S4.

### Cell viability assay

To analyze the effects of MSI2 depletion or compound treatment on cell proliferation, cells were seeded (500 cells/well) in 96-well cell culture plates in complete media. We used increasing concentrations of compounds to calculate IC_50_ values for each cell line. After 72 hours incubation with compounds, we added CellTiter-Blue® reagent (Promega, Fitchburg, WI) to cells and incubated them for 2 hours at 37 ºC. After that we recorded fluorescence (560_Ex_/590_Em_). Data was analyzed using GraphPad Prism software.

### Clonogenic assay

Murine KP, KPM2 cells and human A549 cells (500 cells per well) and H441 cells (2000 cells per well) were plated in 12-well plates and incubated in complete media. After 24hours compounds were added, after which cells were incubated for 7 days. Cells were fixed in 10% acetic acid/ 10% methanol solution and stained with 0.5% (w/v) crystal violet as previously described(37). A colony was defined as consisting of >50 cells and counted digitally using ImageJ software as described previously(38).

### Cell cycle analysis

Cells were seeded in 6 well plates for 72 hours. Cells were washed with cold PBS twice and dissociated with an EDTA-free trypsin solution. Cells were collected and washed twice with cold PBS (cells were centrifuged at 1500 rpm for 5 min after each wash). Cells were fixed with 100 μl of reagent A (#GAS001, Invitrogen) with 4ml absolute methanol for 10 minutes, then incubated with 100 μl reagent B (#GAS002, Invitrogen) for 30 minuets and incubated with 0.1% paraformaldehyde fixative solution for overnight storage at 2–8ºC in the dark. FxCycle™ Violet (#F10347, Invitrogen) were added to stain the cells at 2–8ºC for 1 hour, after that cells were analyzed by flow cytometry.

### RNA-IP assays

RNA was immunoprecipitated from cell lysates (2 × 10^7^ cells per IP) using either a control normal rabbit IgG or rabbit monoclonal anti-MSI2 antibody and the Magna RIP RNA-binding Protein Immunoprecipitation kit (cat#17-700, Millipore, Burlington, MA). Manufacturer’s instructions were followed with the exception that RNeasy MinElute Cleanup kit (cat#74202, Qiagen, Venlo, Netherlands) was used to prepare RNA. Immunoprecipitated RNAs were quantified by quantitative PCR (qPCR) using primers indicated in Supp Table S4, using PTP4A1 as a normalization (positive control) and GAPDH as a negative control.

### RPPA

The murine cell lines were lysed and prepared according to MD Anderson Core Facility instructions(39-41), and RPPA was performed at MD Anderson facility. Heatmap was generated using the GraphPad Prism.

### In silico evaluation of MSI2 binding to mRNAs

Human and murine genome sequences for ATM and other mRNAs from the NCBI (ATM human (NCBI Reference Sequence: NM_000051.4), ATM mouse (NCBI Reference Sequence: NM_007499.3), CHK2 human (NCBI Reference Sequence: NM_001005735.2), CHK2 mouse (NCBI Reference Sequence: NM_001363308.1) were scanned for Musashi binding motifs previously defined by Bennett et al. (15 motifs with highest p values), Wang et al. (8 motifs with highest p values and Nguen et al. (12 motifs with highest p values). (Supp Tables S6, S7, S8).

### Statistical analysis

All statistical analyses, including unpaired two-tailed t-test, ANOVA analysis, Spearman correlation, were performed in GraphPad Prizm 9 (San Diego, CA).

## Results

### Musashi-2 contributes to the incidence of KRAS-P53 driven mouse lung adenocarcinoma and positively regulates lung cancer cell growth

To assess the role of Msi2 in NSCLC development *in vivo*, we generated a new murine model (KPM2) by crossing 129S/Sv-*Kras*^*tm3Tyj*/J^;*Trp53*^*t*m1Brn/J^ (KP)(34) and *Msi2*^*-/-*^ mice (33) (Fig. 1A). In these mice, intratracheal administration of lentivirus expressing Cre recombinase simultaneously activates *Kras*^*G12D*^ and deletes *P53* to initiate tumorigenesis in the lung epithelium, and for those bearing *Msi2*^*-/-*^, Msi2 is concurrently deleted. Induction of Cre in KP and KPM2 mice induced lung tumors in both KP and KPM2 mice after 10 weeks. KPM2 mice had significantly fewer distinguishable independent tumors and a lower total tumor burden (Fig. 1B, Supplementary Fig. S1). In addition, proliferation marker Ki67 is significantly decreased in the KPM2 lung tumors (Fig. 1C). Levels of Msi1 were comparable in the KP and KPM2 lung tumors (Fig. 1C), indicating observed activity was specific to Msi2, rather than involving cross-regulation of Msi1. To define the mechanism underlying the reduced tumor growth phenotype upon Msi2 elimination, we established six murine lung tumor cell lines from KP and KPM2 mice: KP-1, KP-2, KP-3 and KPM2-1, KPM2-2, KPM2-3 respectively (Fig. 1D). Cell growth analysis of murine the lung tumor cell lines using Cell Titer Blue (CTB) and clonogenic assays indicated significantly slower KPM2 cells growth in comparison with KP cells (Fig. 1E and F).

**Figure 1.**
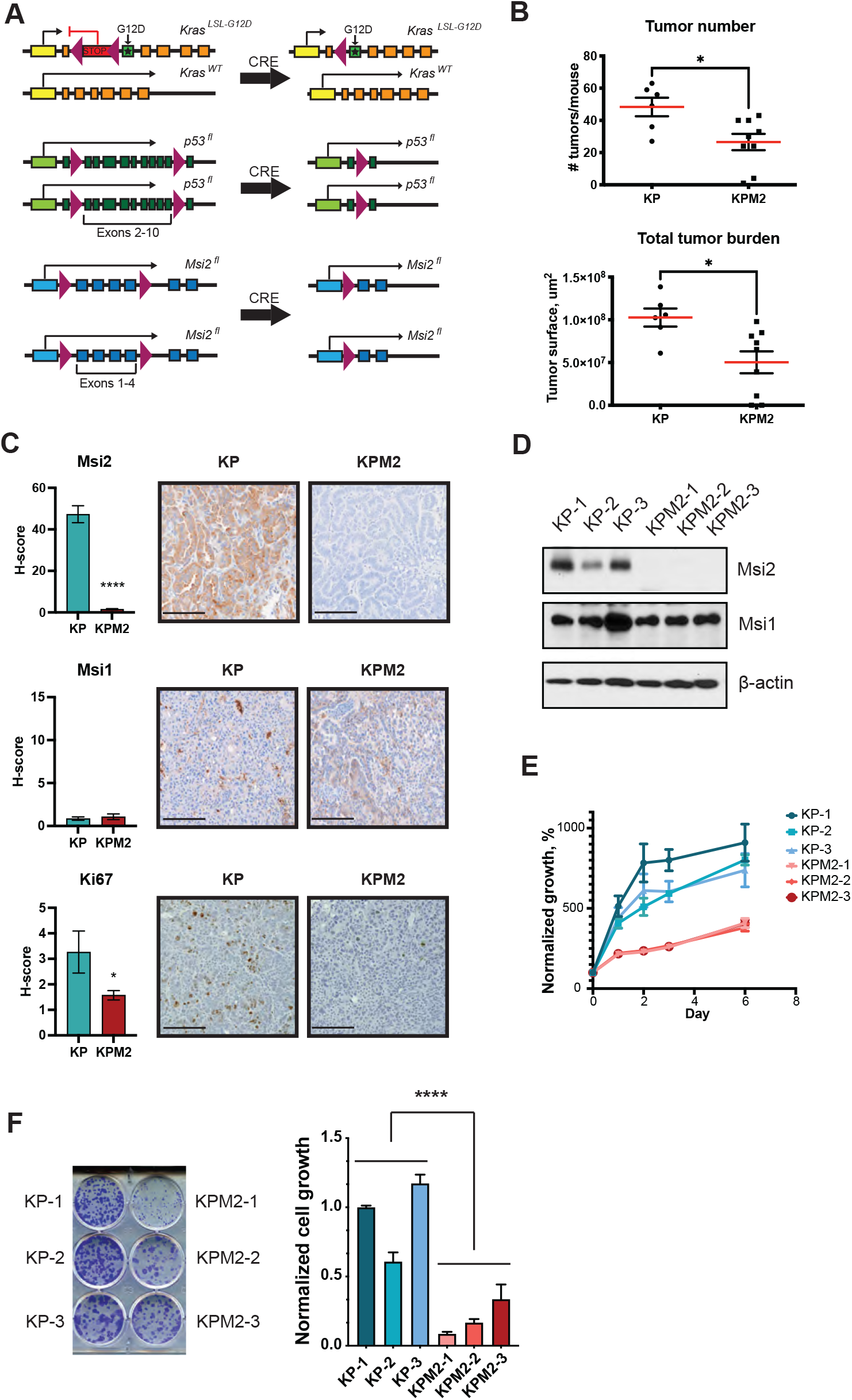
Characterization of novel Musashi 2 deficient murine lung cancer models (KPM2) in comparison to KP. (A) Targeting scheme for Kras-mut, p53 KO and Msi2 KO Cre-inducible mice. We crossed mice with Cre-inducible Msi2 depletion (M2) with the mice that have Cre-inducible Kras mutation (G12D) and p53 depletion (KP) to generate murine model with Cre-inducible Kras mutation (G12D), p53 and Msi2 depletions (KPM2) (B) Quantifications of tumor number and tumor burden per each mouse group. KP groupcontains 6 mice, KPM2 group contains 9 mice. Statistical analysis was performed using a Mann-Whitney test. * p < 0.05. (C) Representative IHC images of Msi2, Msi1, and Ki67 protein expression in murine lung tumors and H-score quantifications. Scale bar 100 μm. (D) Western blots of indicated cell lines. Cell lines were generated from lung tumors derived from KP mice (KP-1, KP-2, KP-3), and from KPM2 mice (KPM2-1, KPM2-2, KPM2-3). (E) Cell growth quantified by Cell Titer Blue (CTB) assay. Data from at least 3 independent repeats and normalized to day 0, day 0 is equal to 100%. (F) Clonogenic assay of murine lung tumor cell lines. Data from at least 3 independent repeats and normalized to KP (average of KP-1, KP-2, KP-3). (C, E, F) Error bars represented by SEM. Statistical analysis was performed using an unpaired two-tailed t-test. *p < 0.05, ****p <0.0001.

### Musashi-2 depletion leads to impaired of DNA damage response signaling and cell cycle in murine and human NSCLC

Since Msi2 KO reduces lung tumor growth in KPM2 *in vitro* and *in vivo*, we performed Reverse Phase Protein Array (RPPA) analysis of KP and KPM2 cell lines (Supplementary Tab S9). As a result, RPPA data indicated that Msi2 depletion leads to decrease of Vegfr2, Smad3, Sox2 (Supplementary Tab S9). That supports previously published papers (22,32,36). In addition, MSI2 may affect DDR signaling in particular, RPPA results suggested that both total Atm and auto phosphorylated Atm (phS1981) protein levels are decreased in cells lacking Msi2 (Fig. 2A, Supplementary Fig. S2A). We then used Western blot analysis of KP and KPM2 cell lines for extended detection DDR signaling proteins (Fig. 2B and C). Our Western blot analysis show ed that phospho-Atm, total Atm and its direct downstream target Chk2 and phospho-Chk2 are all significantly decreased with Msi2 KO despite increase in the level of phospho-H2AX (Fig. 2B). Based on that, we performed western blot analysis to probe all relevant DDR signaling pathways (Fig. 2C). Analysis of Atr downstream signaling showed that phospho-Chk1 (an active form of direct downstream target of Atr) is increased, while Cdc25A protein level was decreased and phospho-Cdc2 increased in Msi2 deficient cells (KPM2), consistent with published data (42,43). Subsequent cell cycle analysis of murine NSCLC cell lines indicates that Msi2 KO leads to a significant decrease percentage of cells in G1/M phase and an increase in percentage of cells in G2/M phase (Fig. 2D, Supplementary Fig. S2B).

**Figure 2.**
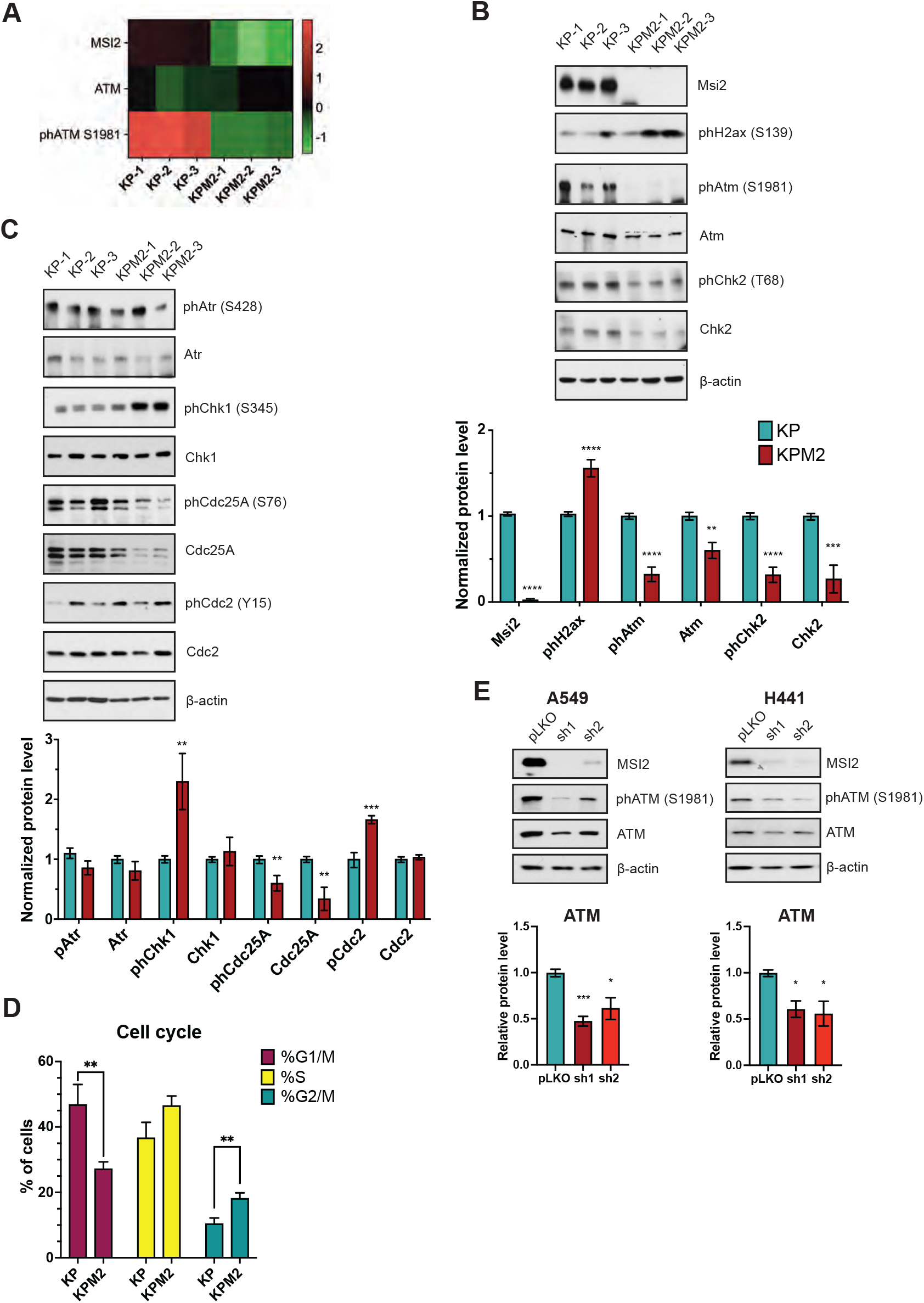
Depletion of MSI2 in murine KP and in human cell lines affects DNA damage response signaling. (A) Heatmap of RPPA result for ATM, phATM protein expression in murine lung cancer cell lines with Msi2 KO (KPM2-1, KPM2-2, KPM2-3) and without Msi2 KO (KP1, KP-2,KP-3). RPPA performed in KP and KPM2 cell lines in two biological repeats. (B,C) Western blotanalysis of indicated murine cell lines. Data normalized to KP (average of KP-1, KP-2, KP-3) and β-actin. Error bars represented by SEM. (D) Cell cycle analysis of murine lung cancer cell lines. KP group contains KP-1, KP-3 cell lines, KPM2 group contains KPM2-1, KPM2-3 cell lines. Quantification was performed from at least three independent experiments by FlowJo software. Error bars represented by SEM. (E) Western blot analysis of human NSCLC cell lines, following depletion by shRNA (sh1, sh2) of MSI2. Negative control includes pLKO transfected cell line. MSI2 depletion was induced by the addition of 1 μg/ml of Doxycycline for 48 h. Data normalized to pLKO and β-actin. Error bars represented by SEM. (B,C,E) Quantification of Western blot data from at least three independent experiments by Image J software. Statistical analysis was performed using an unpaired two-tailed t-test. *p < 0.05, **p <0.01, ***p < 0.001, ****p <0.0001 for all graphs.

To validate MSI2 regulation of ATM in human NSCLC, we used human NSCLC cell lines (A549, H441) with MSI2 doxycycline-inducible knockdown (KD) (sh1, sh2) from our previous papers (21,32) and evaluated phospho-ATM and total ATM levels after MSI2 inducible KD (Fig. 2E). Western blot analysis showed that MSI2 KD leads to significant decrease of both phospho-ATM and total ATM in human NSCLC cell lines. Thus, we conclude that Musashi-2 supports ATM expression in murine and human NSCLC.

Then we used murine lung tumor samples to evaluate the effect of Msi2 KO on DDR signaling (Fig. 3). We performed Western blot analysis of tumor samples from 3 KP and 3 KPM2 mice (Fig. 3A). Our analysis showed dramatic decrease of Atm phospho-Atm and increase of phospho-Chk1, phospho-Cdc2, phospho-H2AX protein levels. Which is consistent with data from murine and human lung cancer cell lines (Fig. 2). Next, IHC analysis of murine tumor samples showed strong positive correlation of Atm with Msi2 (Pearson rank = 0.62) and strong negative correlation of phospho-H2AX with Msi2 (Pearson rank = -0.625) and moderate negative correlation of CyclinB1 with Msi2 (Pearson rank = -0.57) (Fig. 3B,C). Taken together, Musashi-2 supports DDR signaling and cell cycle progression in murine and human lung cancer.

**Figure 3.**
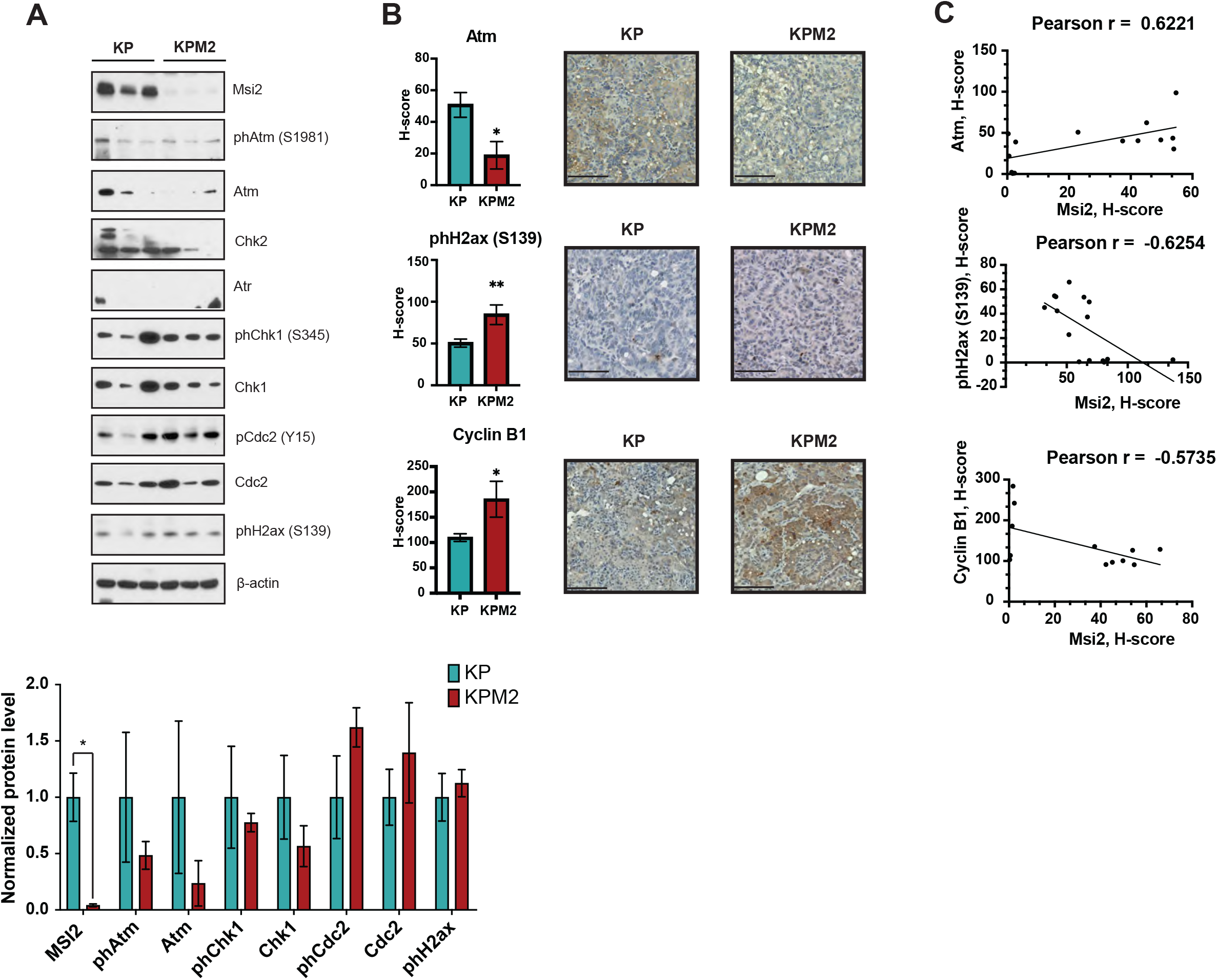
Msi2 depletion results in impared DDR in murine lung tumor samples. (A) Western blot analysis of indicated murine lung tumors. Data normalized to KP (average of 3 biological repeats) and β-actin. Error bars represented by SEM. Quantification of Western blot data from three biological repeats by Image J software. (B) Representative IHC images of Atm, phH2ax (S139), Cyclin B1 protein expression in murine lung tumors and H-score quantifications. Scale bar 100 μm. Statistical analysis was performed using an unpaired two tailed t-test. (C) Correlation analysis of Atm vs Msi2, phH2ax (S139) vs Msi2, Cyclin B1 vs Msi2 protein levels of murine lung cancer samples performed using Pearson correlation coefficient. Number of samples 15. For all graphs, statistical analysis was performed using an unpaired two-tailed t-test. *p < 0.05, **p < 0.01, ***p < 0.001, ****p <0.0001 for all graphs.

### Musashi-2 directly regulates ATM/Atm protein levels in lung cancer

The above data suggests that MSI2 may directly regulate translation of the *ATM/Atm* mRNA. Previously studies (23-25) have determined that MSI2 directly binds consensus sequences with a core motif UAG in the 3’-UTR of target mRNAs, and as a result regulates the stability and translation of target mRNAs. Based on that published data we performed *in silico* analysis and found that *ATM* mRNA has predicted binding sites for Musashi-2 binding (Fig. 4A, Supplementary Tab. S6, S7, S8). To confirm the *in silico* predictions, we did RNA immunoprecipitation assays (RIP) with an MSI2 antibody coupled with RT-qPCR in 3 NSCLC cell lines: murine KP-1, and human A549, H441(Fig. 4A), using previously defined MSI2 target mRNAs (*PTP4A, TGFβR1, SMAD3)(22,33)* as positive controls and *ACTB* and *GAPDH* as negative controls. Antibodies to MSI2 specifically immunoprecipitated the ATM mRNA as efficiently as they did the positive controls (Fig. 4B). In addition, we performed RTqPCR analysis of KP and KPM2 cell lines showed that *Msi2* KO has no effect *Atm* mRNA level (Supplementary Fig. S2). Taken together, we conclude that MSI2 directly regulates ATM/Atm protein level in murine and human NSCLC via direct binding to*ATM/Atm* mRNAs.

**Figure 4.**
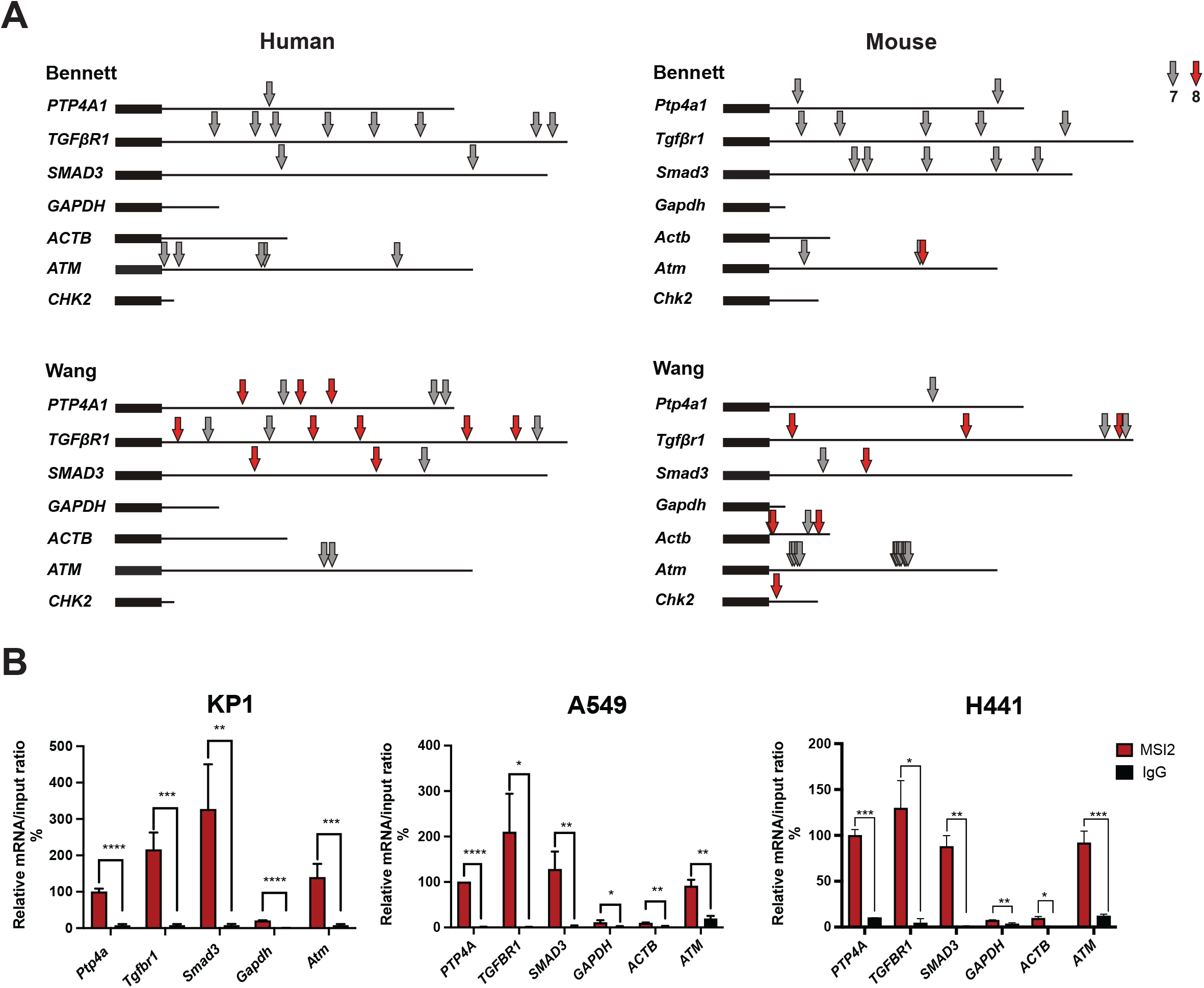
MSI2 directly regulates ATM/Atm protein level via direct binding of ATM/Atm mRNAs. (A) MSI2 consensus binding sites on human and murine mRNA. Location of consensus binding sites for Musashi proteins in the genes, as defined from studies by Bennett et al, and Wang et al. Coding sequences are represented by thick lines; 3’ untranslated regions by a thin line. 7-or 8-bp consensus sequences are indicated by arrows. ATM human (NCBI Reference Sequence: NM_000051.4), Atm mouse (NCBI Reference Sequence: NM_007499.3), CHK2 human (NCBI Reference Sequence: NM_001005735.2), Chk2 mouse (NCBI Reference Sequence: NM_001363308.1), SOX2 human (NCBI Reference Sequence: NM_003106.4), Sox2 mouse NCBI (Reference Sequence: NM_011443.4). (B)Quantification of mRNA immunoprecipitation (RIP) results from assays performed in KP-1, H441 and A549 cell lines lysates using antibodies to MSI2, or IgG (negative control) antibodies, followed by quantitative RT-PCR. Data are normalized to positive control PTP4A1, TGFBR1, and SMAD3 are additional positive controls; GAPDH is a negative control. The data shown reflect the average of three independent RIP experiments. Error bars represented by SEM. Statistical analysis was performed using an unpaired two-tailed t-test.

### Musashi-2 depletion leads to sensitization of murine and human NSCLC to PARP inhibitor treatment

Several clinical trials have recently shown potential efficacy of PARP inhibitors, especially when combined with platinum-based chemotherapy or immunotherapy, although there is lack of clarity on specific biomarkers to select NSCLC patients that might benefit from this approach (13-16,44). In addition, recent publications show that ATM deficiency is associated with higher sensitivity to PARP inhibitor olaparib in NSCLC (18-20). Taken together, we hypothesized that Musashi-2 depletion, which decreased in ATM protein levels, might be associated with sensitization of NSCLC to olaparib treatment. To test this hypothesis, murine NSCLC KP and KPM2 cell lines were treated with olaparib (Fig. 5A and C, Supplementary Fig. S3A). Clonogenic and CTB viability analyses showed significant sensitization in cells with Msi2 KO to olaparib treatment. Olaparib treatment of human NSCLC cell lines also showed significant sensitization of cells with MSI2 inducible knockdown (Fig. 5B and C, Supplementary Fig. S3B).

**Figure 5.**
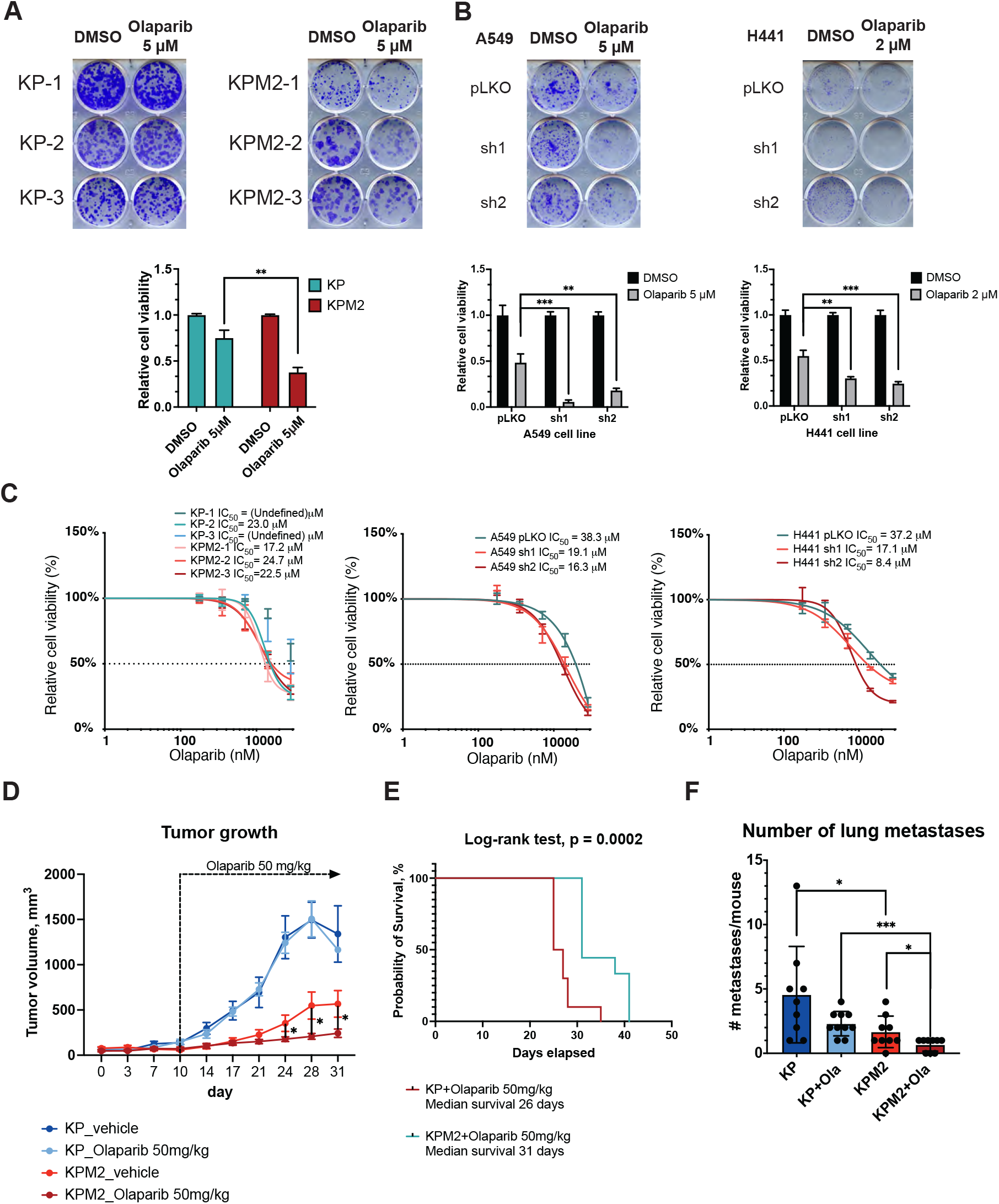
MSI2 depletion leads to olaparib treatment sensitization. (A) Clonogenic assay of murine lung tumor cell lines. Data normalized to DMSO in each group. (B) Clonogenic assay of human NSCLC cell lines, following depletion by shRNA (sh1, sh2) of MSI2. Negative control includes pLKO transfected cells. Data normalized to pLKO. MSI2 depletion was induced by the addition of 1 μg/ml of Doxycycline for 48h. (A, B) Error bars represented by SEM. Statistical analysis was performed using an unpaired two-tailed t-test from at least 3 independent repeats. **p <0.01, ***p < 0.001. (C) Cell viability assay performed by Cell Titer Blue (CTB) assay of indicated cell lines with olaparib treatment. IC_50_ quantifications were performed in GraphPad software from at least 3 independent repeats. (D) Growth curves of subcutaneous allografts of murine lung tumor cells. (E) Kaplan-Meier analysis of mice survival with subcutaneous allografts of murine lung tumor cells with olaparib treatment. (F) Number of lung metastases of subcutaneous allografts of murine lung tumor cells after 31 days of growth. (D, F) Mice were treated with vehicle or olaparib 50mg/kg for 21 days. N=9/group. Statistical analysis was performed using an unpaired two-tailed t-test, *p<0.05, ***p < 0.001.

Then we performed allograft analysis to evaluate the effect of Msi2 KO on olaparib treatment *in vivo*. We used KP-1 and KPM2-2 cell lines for subcutaneous injection. When tumor volumes reached 100 mm^3^, animals were dosed for 21 days with either olaparib (50 mg/kg, 5 days a week) or vechicle. Consistent with the *in vitro* experiments, Msi2 KO leads to tumor growth decrease and olaparib treatment sensitization (Fig. 4D). In addition, overall survival of mice with olaparib treatment was longer with Msi2 KO (Fig. 4E). Moreover, using the H&E staining of murine lungs after 31 days of tumor growth we showed that Msi2 KO significantly decreases number of lung metastasis (Fig. 5F, Supplementary figure S4A), which is consistent with our prior Musashi studies in other lung murine models (22).

Finally, we performed in vitro treatments with platinum-based chemotherapy drug cisplatin. Murine lung tumor cell lines with Msi2 KO have significant sensitization to cisplatin treatment in comparison to KP cell lines (Fig. 3D and E). In contrast, MSI2 KD in human NSCLC cell lines leads to slightly lower sensitivity to cisplatin treatment by CTB viability or no effect in clonogenic assay (Supplementary Fig. S3C and D). Taken together, we conclude that Musashi-2 depletion leads to sensitization of both murine and human NSCLC to PARP inhibitor treatment and to sensitization to platinum-based chemotherapy only in KP murine model, but not in human NSCLC.

## Discussion

In this study, we established new transgenic mouse model by deleting *Msi2* post lung-specific KP tumors induction (KPM2) and demonstrated that *Kras/Trp53*-dependent NSCLC in the absence of Msi2 results in less aggressive tumorigenesis (Fig. 1B). These results inform that Musashi-2 has oncogenic and growth promoting properties during lung tumorigenesis, which is also confirmed by other published papers(21,22,24,30,31). The novelty of our study is that Musashi-2 regulates DDR signaling, in particular via controlling ATM protein level, in lung cancer.

Lung cancer cell lines derived from KPM2 mice demonstrate decreased proliferation and clonogenic survival (Fig. 1E and F). This further confirms the oncogenic role of Msi2 in NSCLC, and suggests it is one of the proteins that are upregulated early during lung carcinogenesis and that sustained Msi2 expression contributes to NSCLC aggressiveness via regulation of downstream signaling. Results of RPPA analysis support previously published papers (22,32,36) and indicated that Msi2 depletion leads to decrease of Vegfr2, Smad3, Sox2 (Supplementary Tab S9). Also, RPPA analysis and its validation showed a new effect of Msi2 deletion in murine tumors. We found that Msi2 deletion has led to increased phospho-H2ax and loss of total and phospho-Atm and total and phospho-Chk2 and G2/M cell cycle arrest and decreased cell proliferation. But these results are somewhat unexpected,because the activation of DDR signaling components leads to cell cycle arrest and decreased cell proliferation(9). Since ATM is a major kinase phosphorylating H2A, and reduction of ATM should have led to loss of phospho-H2Ax. We suspect, paradoxical increase in phospho-H2Ax is mediated by another kinase, possibly DNA-PKc(45).

It is well known that DDR signaling is divided in two parts: ATM-dependent signaling and ATR-dependent signaling, which share substrate specificity, that is they preferentially phosphorylate serine or threonine residues followed by glutamine(11). Based on that, evaluation of ATR-dependent signaling indicated increased phoshpo-Chk1, decreased total Cdc25A, increased phospho-Cdc2 (Y15) (Fig. 2C), which confirms impaired cell cycle and decreased cell proliferation (Fig. 2D, Fig. 1E and F). Also, IHC analysis and of murine lung tumors indicates strong positive correlation Msi2 with Atm and strong negative correlation Msi2 with phospho-H2ax and Msi2 with CyclinB1 (Fig. 3B) Otherwise, human NSCLC cell lines showed depletion of total and phospho-ATM with MSI2 KD (Fig. 2E). Thus, we confirm that decrease of Atm/ATM associated with Musashi-2 depletion and leads to DDR signaling disruption, accumulation of DNA breaks, impaired cell cycle and inhibition of cell growth. This is an exciting new finding, which shows previously unexpected role of MSI2 in control of DNA damage response in lung cancer cells.

As Musashi-2 is known as RNA-binding protein, we showed that Musashi-2 directly binds and regulates mRNA translation of *ATM/Atm* using *in silico* analysis and RNA-IP analysis (Fig. 4A,B). That result explains why NSCLC models become more sensitive to olaparib with depletion of Musashi-2. Therefore, our results expand knowledge about the role of Musashi-2 in cancer. Published papers by our group and other groups have already showed that Musashi-2 directly regulates EGFR, TGFbR1, SMAD3, PTEN etc.(21,22,24,31), and ATM regulation is another important one. IHC analysis of murine lung tumors and TMA analysis of human lung tumor samples show strong positive correlation between ATM/Atm and MSI2/Msi2 protein levels which may suggest use of Musashi-2 and/or ATM protein levels as predictive biomarker of response to DDR therapy in NSCLC.

Importantly, we also show that deletion of Msi2 in murine, or depletion of MSI2 in human models leads to sensitization of lung cancer cells to PARP inhibitor treatment *in vitro* (Fig. 5A-C, Supplementary Fig. S3A,B) and *in vivo* (Fig. 5D). In addition, mice injected with Msi2 depleted murine cells show better overall survival (Fig. 5E) and lower amount of lung metastases (Fig. 5F, Supplementary Fig. S4A)These effects are likely mediated by Atm/ATM loss as a result Musashi-2 depletion, as these results are consistent with previously published papers describing PARP inhibitor sensitization of NSCLC cells due to ATM loss(18-20). In addition, we show that Msi2 depletion in murine lung cancer cells, but not in human cell lines, leads to sensitization to cisplatin (Supplementary Fig. S3C-F). It can be possibly related to *TP53* status in those models, because while murine cell lines have*Tp53* depletion, H441 cell line has *TP53* mutation (473G>T) and A549 cell line has *TP53* wild-type. Moreover, published papers show that both wild-type and some mutant p53 forms could mediate cisplatin resistance(46,47).

Finally, because of the absence of easily druggable domains in noncatalytic RNA-binding proteins, it is challenging to develop effective small molecule inhibitors targeting Musashi proteins. A structurally unrelated neural growth factor (NGF) inhibitor developed by Roche, Ro 08-2750(48,49), has recently been found to inhibit Msi1/Msi2 in vitro and in vivo(50). We have recently developed a first in class Musashi inhibitor, which effectively interferes with RNA Recognition Motif 1 (RRM1) binding of Musashi proteins to its targets(51). Further refinement of these inhibitors, including generation of more potent compounds or degraders of Musashi proteins, is warranted, to allow development of new cancer therapeutic approaches.

## Acknowledgements

This work was supported by the Northwestern University Cancer Center Support Grant (NCI CA060553). The Functional Proteomics RPPA Core is supported by MD Anderson Cancer Center Support Grant # 5 P30 CA016672-40. This work and the authors were supported by NIH R01 grant CA218802 (to Y.B.); a Translational Bridge Award from Northwestern University, number 2022-001 (to Y.B.); NCI Core Grant P30 CA006927 (to the Fox Chase Cancer Center), NCI R21 grant CA263362 (to P.M.); by DOD CA201045 / W81XWH2110487 and by the William Wikoff Smith Charitable Trust (to E.G.)

---

**Figure S1.**
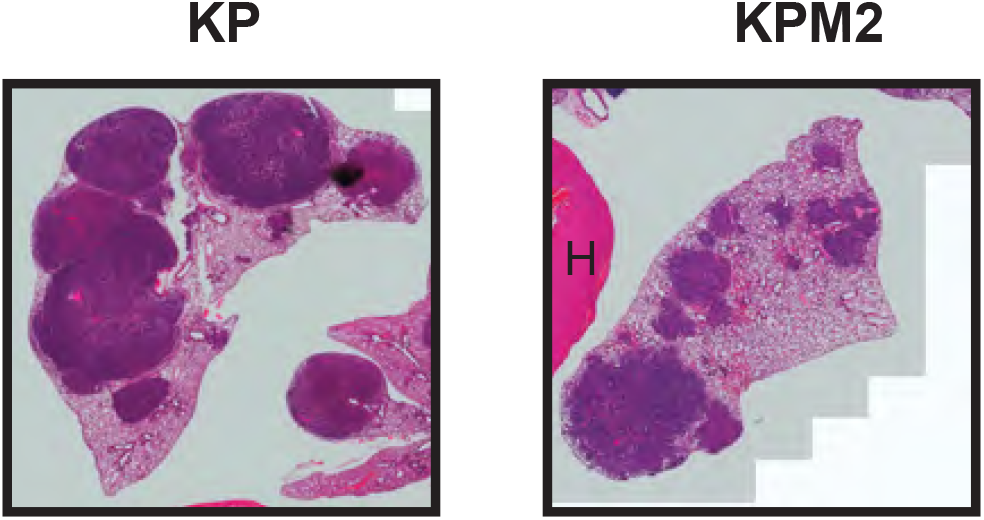
Hematoxylin and eosin-stained lung and heart (H) tissues.

**Supplementary figure S2.**
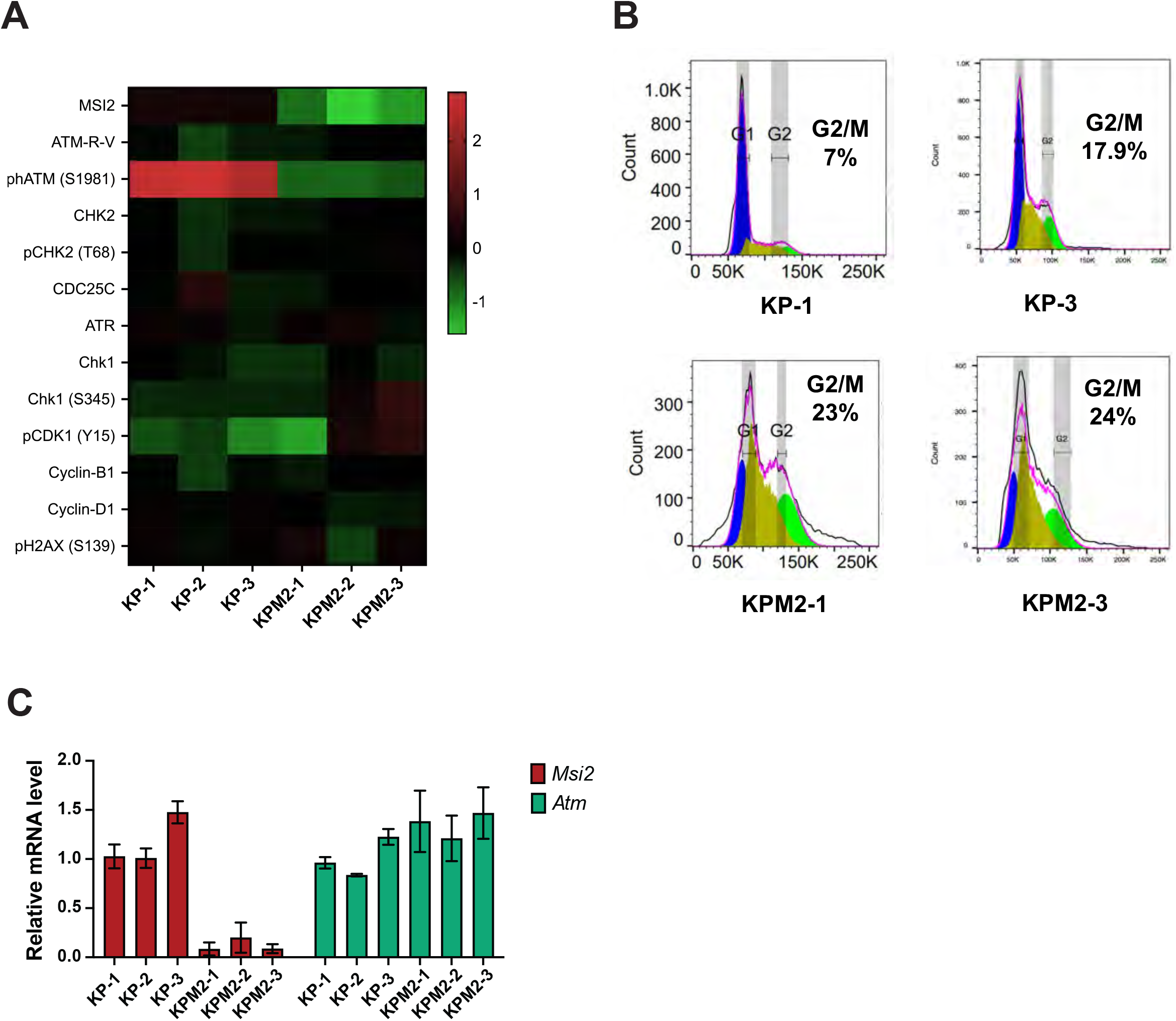
(A) Heatmap of RPPA result for DNA damage response signaling proteins expression in murine lung cancer cell lines with Msi2 KO (KPM2-1, KPM2-2, KPM2-3) and without (KP1, KP-2, KP-3). RPPA performed in KP and KPM2 cell lines in 3 biological repeats and 2 technical repeats. (B) Representative graphs of cell cycle analysis of murine lung cancer cell lines. Quantifications were performed in FlowJo software. (C) RTqPCR analysis of Msi2 and Atm mRNA in murine lung cancer cell lines.

**Supplementary figure S3.**
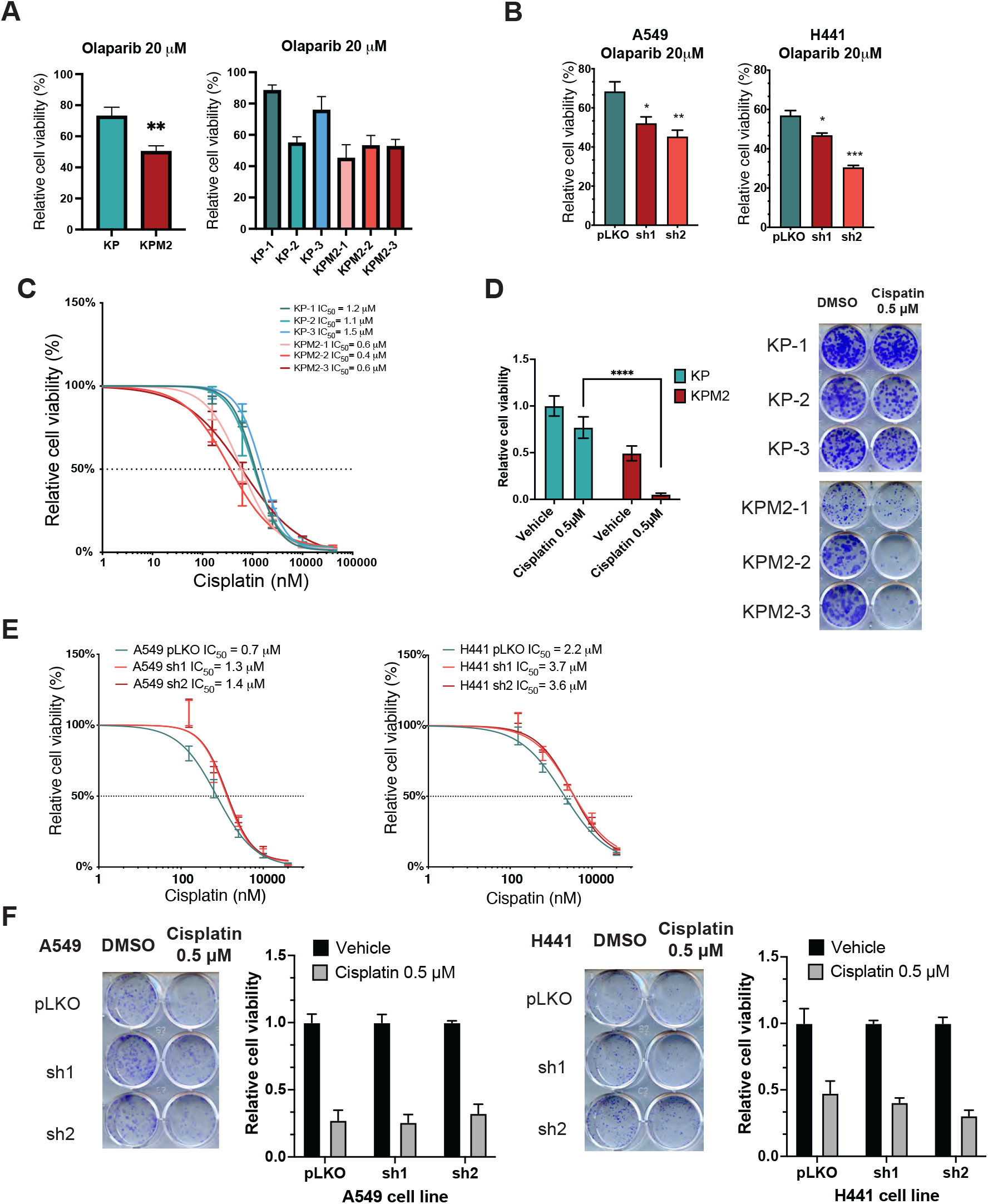
(A) Viability analysis with 20 μM olaparib treatment for 72 hours of indicated murine lung tumor cell lines. KPM2 is group of cell lines with Msi2 KO (KPM2-1, KPM2-2, KPM2-3), KP is group without Msi2 KO (KP1, KP-2, KP-3). Data were extracted from IC50 analysis (Figure 3D). (B) Viability analysis with 20 μM olaparib treatment for 72 hours of indicated human NSCLC cell lines, following depletion by shRNA (sh1, sh2) of MSI2. Negative control includes pLKO. MSI2 depletion was induced by the addition of 1 μg/ml of Doxycycline for 48 h. (C) Cell viability assay performed by Cell Titer Blue (CTB) assay of indicated cell lines with Cisplatin treatment. IC_50_ quantifications were performed in GraphPad software. (D) Clonogenic assay of murine tumor cell lines. Data normalized to Vehicle in each group. (E) Cell viability assay performed by Cell Titer Blue (CTB) assay of human NSCLC cell lines with Cisplatin treatment. IC_50_ quantifications were performed in GraphPad software from at least 3 independent repeats. (F) Clonogenic assay of human NSCLC cell lines, following depletion by shRNA (sh1, sh2) of MSI2. Negative control includes pLKO transfected cells. Data normalized to pLKO. MSI2 depletion was induced by the addition of 1 μg/ml of Doxycycline for 48h. Statistical analysis was performed using an unpaired two tailed t-test. For all graphs: error bars represented by SEM, data from at least 3 independent repeats, *p < 0.05, **p < 0.01, ***p < 0.001.

**Supplementary figure S4.**
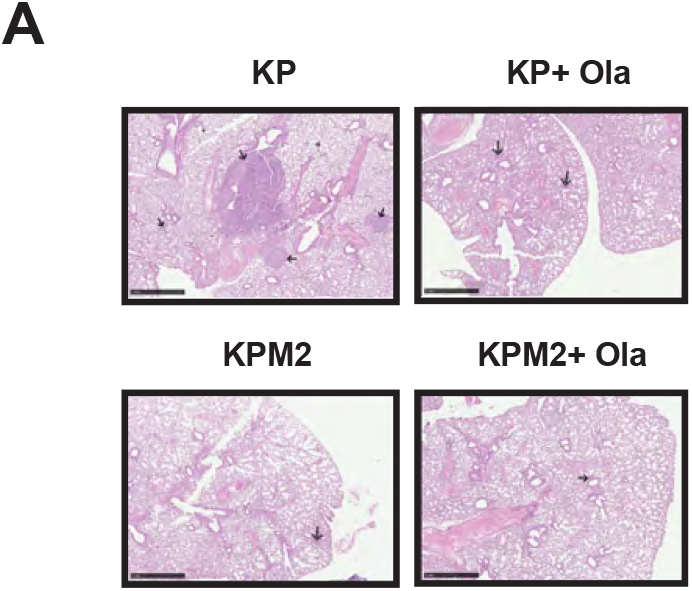
A) Hematoxylin and eosin-stained murine lung of subcutaneous allografts of murine lung tumor cells after 31 days of growth. Mice were treated with vehicle or olaparib 50mg/kg for 21 days. N=9/group. Lung metastases are marked with arrows. Scale bar 1mm.

## Notes

### Competing Interest Statement

The authors have declared no competing interest.

